# The *Marchantia* pangenome reveals ancient mechanisms of plant adaptation to the environment

**DOI:** 10.1101/2023.10.27.564390

**Authors:** Chloé Beaulieu, Cyril Libourel, Duchesse Lacourt Mbadinga Zamar, Karima El Mahboubi, David J. Hoey, Jean Keller, Camille Girou, Helene San Clemente, Issa Diop, Emilie Amblard, Anthony Théron, Stéphane Cauet, Nathalie Rodde, Sabine Zachgo, Wiebke Halpape, Anja Meierhenrich, Bianca Laker, Andrea Brautigam, George RL Greiff, The SLCU Outreach Consortium, Peter Szovenyi, Shifeng Cheng, Yasuhiro Tanizawa, James H. Leebens-Mack, Jeremy Schmutz, Jenel Webber, Jane Grimwood, Christophe Jacquet, Christophe Dunand, Jessica M. Nelson, Fabrice Roux, Hervé Philippe, Sebastian Schornack, Maxime Bonhomme, Pierre-Marc Delaux

**Author notes:** Unité de Recherche Physiologie, Pathologie, et Génétique Végétales (PPGV), INP PURPAN, Université de Toulouse, Toulouse, France. Department of Insect Symbiosis, Max Planck Institute for Chemical Ecology, 07745 Jena, Germany (Jean Keller). University of Bristol, 24 Tyndall Avenue, Bristol, BS8 1TQ, UK. Equal contribution (Chloé Beaulieu, Cyril Libourel).

## Abstract

Plant adaptation to a terrestrial life 450 million years ago played a major role in the evolution of life on Earth. This shift from an aquatic environment has been mostly studied by focusing on flowering plants. Here, we gathered a collection of 133 accessions of the non-vascular plants *Marchantia polymorpha* and studied its intraspecific diversity using selection signature analyses, genome-environment association study and a gene-centered pangenome. We identified adaptive features shared with flowering plants, such as peroxidases or nucleotide-binding and leucine-rich repeat (NLR), which likely played a role in the adaptation of the first land plants to the terrestrial habitat. The *M. polymorpha* pangenome also harbored lineage-specific accessory genes absent from seed plants. We conclude that different land plants lineages still share many elements from the genetic toolkit evolved by their most recent common ancestor to adapt to the terrestrial habitat, refined by lineage specific polymorphisms and gene family evolutions.

## Introduction

The transition of plants from an aquatic to a terrestrial habitat 450 million years ago was one of the major revolutions in Earth history ^1^. The first land plants faced new environmental challenges ranging from drought and UV radiation, nutrient scarcity and the presence of new types of parasitic microorganisms ^2^.

Following this colonization event, two main plant lineages had already diversified 400 million years ago, as supported by molecular clock analyses and the fossil record ^3^. The extant vascular plants, or tracheophytes, encompass lycophytes, ferns, gymnosperms and all flowering plants. The other group, known as bryophytes or non-vascular plants encompasses hornworts, mosses and liverworts.

The long-lasting and successful colonization of terrestrial habitats by plants is the result of adaptations to numerous environmental challenges. Key adaptations evolved in the latest common ancestor of land plants, such as mutualistic associations with symbiotic fungi ^4^, stomata ^5^ or the cuticle ^6^. In addition, every plant lineage continuously adapted to new environmental niches, leading to the distribution of plants in most ecosystems on Earth. Genetic mechanisms underlying adaptation of plants to various biotic and abiotic conditions, such as cold ^7^, pathogens ^8–10^ or symbiotic associations ^4^ have been recently described through phylogenomic approaches. In addition, the response of individual plants to environmental factors has been very well studied in vascular plants and in bryophytes, leading to the discovery of transcriptomic and physiological responses to stresses such as high temperature, drought and salt stress that are largely conserved across land plants 11,12.

Evolutionary divergence or conservation of traits among species emerge from neutral and adaptive processes acting within species. In plants, the genetic bases of population adaptation to environmental constraints such as climate variation have been recently addressed by Genome-Environment Association (GEA) studies in crops and in a selection of non-domesticated model angiosperm species 13. These led to the discovery of a multitude of genetic variants ranging from Single Nucleotide Polymorphism (SNP) 14 to structural and gene presence-absence variations when species pangenomes were used 15,16 demonstrating the complexity and diversity of the genetic bases of adaptation in angiosperms. This focus on angiosperms does not allow to capture the diversity of adaptive strategies evolved by other land plant lineages over the last 450 million years 17.

Among the bryophytes, the liverwort *Marchantia polymorpha* has become a go-to model species to address fundamental questions in plant evo-devo and general biology, such as the colonization of land by plants ^18^ or the mechanisms defining cell polarity in eukaryotes ^19^. In the present study, we collected 133 wild accessions of the liverwort *Marchantia polymorpha*, covering the three subspecies (*M. polymorpha* ssp*. ruderalis, M. polymorpha* ssp. *montivagans* and *M. polymorpha* ssp. *polymorpha*) and resequenced their nuclear genome. This dataset was augmented with two long-read genomes for one European and one North American accession. This sequencing effort provides the first genomic and *in vitro* resource for intra-specific diversity in a bryophyte. Pangenomes for the *Marchantia* genus and for the *M. polymorpha* ssp. *ruderalis* were built. Exploring the SNP landscape by GEA and selective pressures analyses, as well as gene presence-absence variation, revealed the strategies deployed by *M. polymorpha* to adapt to diverse environments. Overall, our findings suggest that environmental adaptation in *M. polymorpha* is achieved by ancestral mechanisms as well as lineage-specific gene polymorphisms and family expansions, and horizontal gene transfer.

## Results

### *Marchantia polymorpha* intraspecific diversity resources

*M. polymorpha* encompasses three subspecies, *M. polymorpha* ssp. *ruderalis, M. polymorpha* ssp. *montivagans* and *M. polymorpha* ssp. *polymorpha* which are distributed in the whole northern hemisphere, excluding the arctic regions ^20^. For more than two decades, two accessions isolated in Japan, Takaragaike-1 (Tak-1, male) and Tak-2 (female), have been widely used by the community, and the Tak-1 nuclear genome, composed of eight autosomes and two sexual chromosomes, was sequenced in 2017 ^18^. To maximize the intraspecific diversity, we collected a total of 133 additional accessions from four main geographic areas between 2013 and 2022: the south of France, the UK, a transect between Switzerland and the north of Sweden, and the USA (Figure 1a, Table S1). The collection sites ranged from sidewalks to soil nearby ponds, and from sea level to an altitude of 1124 m. Most of the UK accessions were collected in the framework of a citizen science project ^21^. To ensure long-term conservation and future distribution of this collection, gemmae were collected from a single individual for each accession and propagated *in vitro* following sterilization. This collection includes 103 accessions from *M. polymorpha* ssp. *ruderalis*, sixteen from *M. polymorpha* ssp. *montivagans* and fourteen from *M. polymorpha* ssp. *polymorpha*. All accessions were sequenced by Illumina with an expected mean genome coverage of 110X (Table S1). Cleaned scaffolds were *de novo* assembled (N50average=49.8 kbp, N50min=2kbp and N50max=78.6 kbp, Table S2) and annotated, leading to an average of 19 705 genes predicted per accession. The average BUSCO completeness of these predictions reached 88.5% (Table S2). In addition, the *M. polymorpha* ssp. *ruderalis* accessions CA, collected in the USA, and BoGa, from Osnabrück in Germany, were sequenced using long-read technologies. These two accessions were chosen to maximize geographic (America and Europe) and genetic diversities (see below). The long-read sequencing of CA and BoGa led to genome assemblies composed of 109 contigs (N50=5.991Mb and L50=12) and to a chromosome-level assembly with nine chromosome sequences and two organellar sequences (N50=26.28Mbp and L50=4), respectively. Structural annotation predicted 21 327 / 21 364 genes, corresponding to a BUSCO completeness of 94.4% / 94.1%, respectively in CA and BoGa. The CA genome assembly and the gene annotations for the 133 accessions are available on MarpolBase (https://marchantia.info). The BoGa genome assembly is available under https://doi.org/10.4119/unibi/2982437. Altogether, we provide here the first genomic and physical collection covering the intra-specific diversity of a bryophyte.

**Figure 1.**
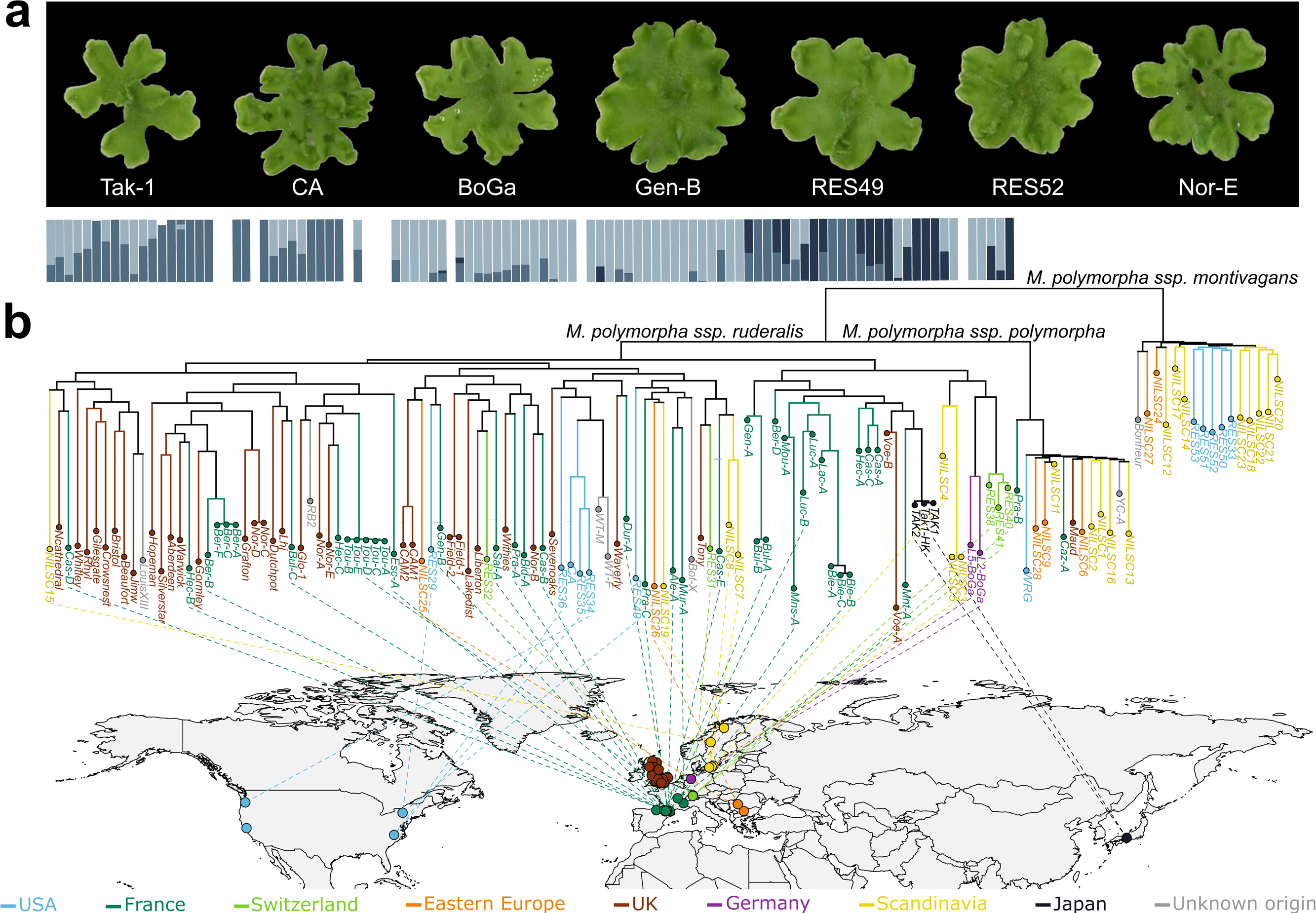
*Marchantia polymorpha* collection, phylogenetic tree and population structure. a) Pictures of representative *Marchantia polymorpha* accessions. Photos were cropped and pasted on a black background. Tak-1, CA, BoGa, Gen-B, RES49 and Nor-E belong to *M. polymorpha* ssp *ruderalis*. RES52 belongs to *M. polymorpha* ssp *montivagans*. b) Phylogenetic tree of the *Marchantia polymorpha* subspecies complex, comprising 104 *M. polymorpha* ssp *ruderalis*, 16 *M. polymorpha* ssp *montivagans* and 13 *M. polymorpha* ssp *polymorpha*. The phylogenetic tree was computed using a dataset of 107 934 pruned SNPs, using the SYM+ASC+R6 substitution model and has a log-likelihood of −1531777.9201. Dots on the world map indicate the sampling location, and colors represent country or geographic regions of origin. The colored branches on the tree also correspond to the geographic origin. Barplots on the top of tree indicate the assignment probability of each *M. polymorpha* accession to each of the three main genetic groups (colored by a shaded blue/grey) identified by the Bayesian clustering algorithm implemented in the fast STRUCTURE software.

### *Marchantia polymorpha* population structure

To characterize intraspecific diversity, reads from all accessions were mapped on the reference Tak-1 genome (∼220 Mb) and SNPs were computed. We identified 12 519 663 SNPs in the collection including all three *M. polymorpha* subspecies. Among these SNPs, 5 414 844 (∼ 1 SNP each 40 bp on average) segregated in *M. polymorpha* ssp. *ruderalis*. The SNP data mapped on the Tak-1 assembly can be explored on MarpolBase (https://marchantia.info).

We reconstructed the phylogeny of *M. polymorpha*’s accessions using a dataset of 107 934 selected SNPs, resolving three main clades corresponding to the three *M. polymorpha* subspecies (Figure 1b). Although a denser sampling effort in the UK and France increased phylogenetic clustering in these regions, we observed a weak correlation between genetic and geographical distances among accessions within the *M. polymorpha* ssp. *ruderalis* clade (Mantel statistic r = 0.087, p = 0.074), as well as long terminal branches, suggesting substantial gene flow and recombination events across its broad geographical range. Bayesian clustering population structure analyses within *M. polymorpha* ssp. *ruderalis* identified three main genetic groups, though some unclear individual assignment probabilities also suggested gene flow events among groups (Figure 1b).

Altogether, these results support *M. polymorpha* as a subspecies complex ^22^ undergoing gene flow throughout a broad geographical range, as shown in *M. polymorpha* ssp. *ruderalis*.

### Conserved selective pressures across the land plant lineage

Using the SNP data for *M. polymorpha* ssp. *ruderalis,* we determined the genome-wide population scaled mutation rate (Watterson’s theta estimator, *θ_w_* ^23^), which was similar to estimates in populations of angiosperms species with comparable sample sizes ^24, 25^. The average value of gene-based Tajima’s *D* and Zheng’s *E* statistics were negative (*D =* −0.356, *E =* −0.317), indicating a genome-wide excess of low-frequency variants (Figure S1). This signature suggests demographic expansion (perhaps accelerated by recent horticultural trade ^26^) with low population substructure in *M. polymorpha* ssp. *ruderalis*, in agreement with the phylogenetic and population structure analyses.

Among the main SNPs categories, 80% were in intergenic regions. In genic regions, SNPs were similarly frequent in intronic (9%) and in non-intronic (11%) regions – that is exons and UTR regions – (Figure 2a). The genome-wide allele frequency spectrum based on these SNP categories was clearly L-shaped as expected under neutral evolution. We observed an excess of rare missense, 5’ UTR, 3’ UTR and the gain of premature start codon in 5’ UTR, relative to SNPs located in intergenic regions, introns, or synonymous SNPs. This distribution most probably reflects an excess of purifying selection on the formers (Figure 2a). Loss-of-function SNPs were predominantly leading to the gain (51%) and less frequently to the loss (25%) of stop codons (Figure 2b). The genome-wide allele frequency spectrum of such SNPs was also L-shaped, with an excess of rare stop gains and splice acceptor SNPs. This reflects an excess of purifying selection against variants leading to molecular phenotypes associated with truncated proteins and retained introns (Figure 2b).

**Figure 2.**
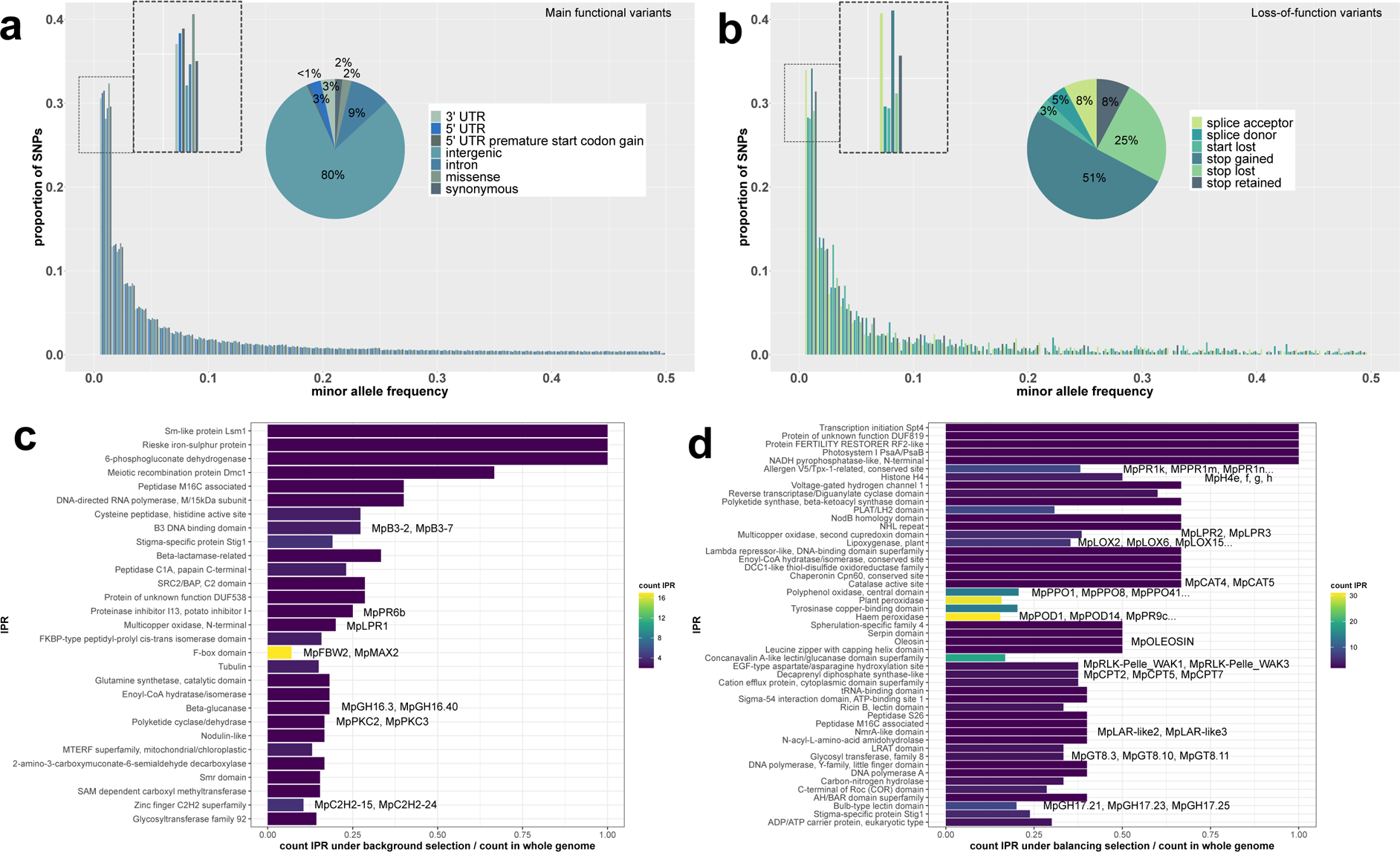
Genome-Wide SNP distribution, gene selection patterns and functional enrichment analysis. In (a), a total of 5 439 674 SNPs (4 363 099 intergenic, 509 964 in introns, 115 218 missense, 88 163 synonymous, 183 858 3’_UTR, 154 821 5’_UTR and 24 551 5’_UTR_premature_start_codon_gain – i.e. a variant in 5’ UTR region producing a three base sequence that can be a START codon –) were used to construct the allele frequency spectrum. In (b), a total of 7 823 SNPs (4 010 stop gained, 1 948 stop lost, 608 stop retained, 606 splice acceptor, 385 splice donor and 266 start lost) were used to construct the allele frequency spectrum. All SNPs in (a) and (b) are distributed on the 8 autosomes and sexual chromosome U or V. In (c), functional enrichment analysis of a list of 777 genes putatively under background selection (top 10% genes of Zheng’s *E* negative value). In (d), functional enrichment analysis of a list of 1 814 genes putatively under balancing selection (top 10% genes of Tajima’s *D* positive values). In (c) and (d), IPR terms are ordered from top to bottom according to decreasing significance (FDR q-value < 0.05), together with some associated known gene names.

To investigate selective pressures on *M. polymorpha* ssp. *ruderalis* genes, we used a dataset of 5 414 844 SNPs for Tajima’s *D* calculation and a dataset of 1 344 013 SNPs with known ancestral/derived allele state, for *H* and *E* calculation. These statistics contrast low, intermediate and high frequency alleles (derived alleles in the case of *H* and *E* statistics). Out of 18 620 genes, we extracted a list of 3 965 genes with the most pronounced selection signatures (777 under background selection, 1 374 under soft/hard selective sweep, and 1 814 under balancing selection, see Method) according to *D*, *H* and *E* statistics (Table S3). We cross-referenced this dataset with similar selection signature analyses in angiosperm species by computing the same statistics on available population genomic SNP datasets in *Arabidopsis thaliana* and *Medicago truncatula* (Table S4) and used orthology relationships to compare selection signatures across species (Table S5). To account for the deep and complex evolutionary divergence between bryophytes and angiosperms, notably due to several rounds of whole genome duplication, we focused on Hierarchical Orthogroups (HOG) showing in each of the three species at least one gene which is in their top 20% of genes under background, balancing or sweep selection at the intraspecific level (Table S5). Functional enrichment analysis of the list of *M. polymorpha* ssp. *ruderalis*’s genes with pronounced signature of background selection revealed molecular functions with reduced polymorphisms within the *M. polymorpha* ssp. *ruderalis* subspecies (Figure 2c, Table S6). Among them, we identified the Receptor-Like Kinase (RLK) protein of the CrRLKL1L-1 family *MpFERONIA/MpTHESEUS* crucial for *Marchantia* and angiosperm development (Table S5) ^27–29^, the DNA repair proteins Dmc1/Rad51/RecA that are essential for meiotic recombination and highly conserved in eukaryotes ^30^, and the Histone-fold Hap3/NF-YB family protein ^31^.

In *M. polymorpha ssp. ruderalis*, functional enrichment analysis of genes under balancing selection revealed a wide array of functions involving highly polymorphic genes, such as the peroxidases (31 genes) and lipoxygenases (nine genes, Figure 2d, Table S6) also found among genes under balancing selection in *A. thaliana* and *M. truncatula* (Table S5). By contrast, fundamental molecular functions seemed to involve highly polymorphic genes specifically in *M. polymorpha*, such as nucleosome with four histone genes (H4e, H4f, H4g, H4h), or DNA-dependent replication with the Y family of DNA polymerases which replicates and repairs damaged DNA (Figure 2d, Table S6). Finally, enrichment analyses based on genes putatively under selective sweep in *M. polymorpha ssp. ruderalis* showed a wide variety of molecular mechanisms and processes, among which various functions such as transcription factors or glycoside hydrolases family 16 (Figure S2, Table S6).

Analyzing selection signatures in *M. polymorpha* ssp. *ruderalis* and comparing it to two representative angiosperms allowed to identify genes and gene families with seemingly conserved selection regime since the latest common ancestor of land plants, following the parsimony principle. In the future, integrating similar data from green algae will allow determining which subset of these genes contributed to the emergence of the land plants.

### Adaptation to climatic constraints in *Marchantia polymorpha* ssp. *ruderalis*

The genome-wide average recombination rate was estimated at 1.5×10^-9^ crossing-over per base pair per meiosis, ranging from 1.2×10^-9^ to 1.7×10^-9^ across the eight *M. polymorpha* ssp. *ruderalis* autosomes. The one-half linkage disequilibrium (LD) decay ranged from 2.5 (chromosome 8) to 4.7 (chromosome 1) kbp across the eight *M. polymorpha* ssp. *Ruderalis* autosomes, with a genome-wide average of 3.6 kbp. Genetic mapping can be thus implemented through genome-wide association approaches (Figure S3). We performed GEA on 96 accessions using 19 bioclimatic variables (BIO1 to BIO19) related to precipitations and temperature, as well as data on monthly average precipitations, solar radiation, water vapor pressure and altitude (see the list of all detected genomic regions in Table S7). To integrate the correlations among bioclimatic variables, we first ran a principal component analysis on the bioclimatic variables related to (i) temperatures (BIO1 to BIO11) on one side and (ii) precipitation (BIO12 to BIO19) on the other. The first three principal components of temperature and precipitation related variables, explaining 95 and 98.4% of the total variance respectively, were retained to describe the main results of the GEA. As an example, a clear geographical structuring of temperature variation is illustrated by principal components plots of *M. polymorpha* ssp. *ruderalis* individuals (Figure 3a) and contributions of the bioclimatic variables to these principal components (Figure 3b).

**Figure 3.**
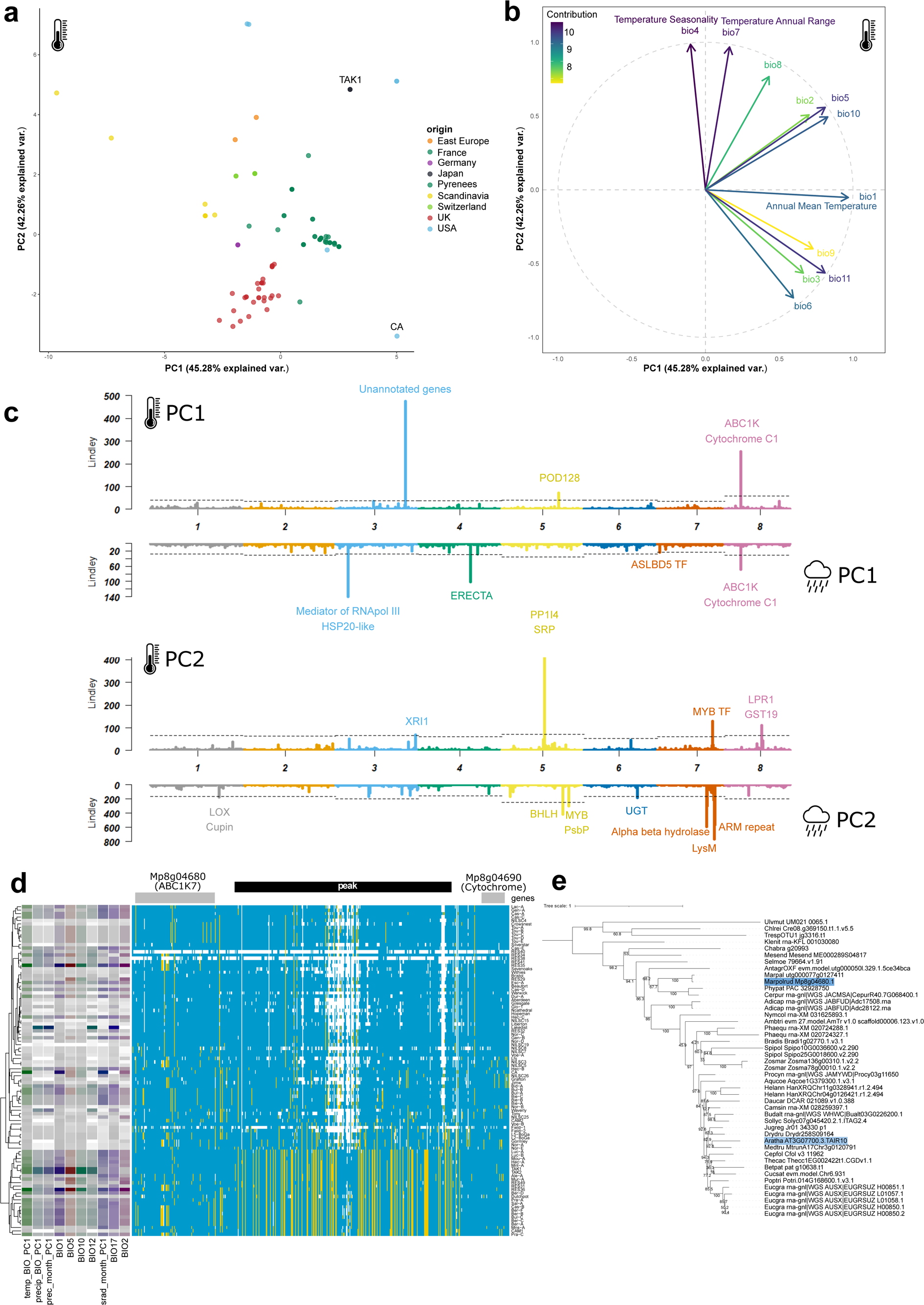
Identification of loci associated with climatic conditions by genome-environment association analyses in *Marchantia polymorpha ssp ruderalis*. (a) Principal components plot of *M. polymorpha* ssp *ruderalis* individuals based on temperature-linked bioclimatic variables (BIO1 to BIO11). (b) Contributions of the temperature-linked bioclimatic variables (BIO1 to BIO11) to the two first principal components. (c) Miami plots of the GEA results on the first components of distinct PCA for 2 categories of bioclimatic variables from the WorldClim2 data: temperature-linked (BIO1 to BIO11) and precipitation-linked variables (BIO12 to BIO19). These Miami plots result from a classical GWAS analysis performed with GEMMA, followed by the use of the local score technique on the SNP p-values, to amplify the signal between SNPs in LD. The dotted lines represent the significance thresholds (resampling thresholds from the local score method). (d) Haplotype block illustration of the genomic region on chromosome 8 associated with both temperature-linked and precipitation-linked variables in *M. polymorpha* ssp *ruderalis*, and containing two protein-coding genes: an ABC1 atypical kinase (Mp8g04680) and a cytochrome c1 protein (Mp8g04690). (e) Phylogenetic tree of the orthologs of the ABC1 gene of *M. polymorpha* ssp *ruderalis* illustrating a direct orthology with the ABC1K7 gene in *A. thaliana.* This tree was computed with the substitution model Q.plant+R5 and has a log-likelihood of −26208.0663.

We identified genomic regions and candidate genes robustly associated with both temperature and precipitation variation across the range of *M. polymorpha* ssp. *ruderalis* (Figure 3c). Among the identified loci was a region on chromosome 8 with a cytochrome c1 protein (Mp8g04690) and an ABC1 atypical kinase (Mp8g04680). In this locus the minor haplotype, present in one quarter of the accessions (24/96), is associated with high values of temperature and precipitation (Figure 3d). The ABC1K gene of this locus is in the top 16% of *M. polymorpha* ssp. *ruderalis* genes under balancing selection, while we did not observe the same tendency in its counterparts in *A. thaliana* and *M. truncatula* (genes are not in the top 20% of genes under balancing selection) (Table S3, S5). ABC1K is orthologous to the ABC1K7 gene in *A. thaliana* (Figure 3e), which acts in concert with ABC1K8 to regulate response to the stress hormone ABA to moderate oxidative stress response ^32^. ABA has been proven to have a role in desiccation tolerance in *M. polymorpha* ^33^, which could explain the link between the ABC1K gene and fluctuations in temperature and precipitations. Other loci were detected as more specifically associated with temperature, such as the peroxidase MpPOD128 (Mp5g17500). This gene belongs to a group of 25 *M. polymorpha* peroxidases, four of which are located in a gene cluster with the candidate, originating from a *M. polymorpha* specific duplication (according to the RedoxiBase ^34^). Finally, several loci were associated only with precipitation variation, such as lipoxygenase MpLOX5 (Mp1g21930) or the region close to the ERECTA (Mp4g14620) gene, the pro-ortholog of three *A. thaliana* genes, ERECTA (AT2G26330), ERECTA like 1 (AT5G62230) and ERECTA like 2 (AT5G07180), which are LRR-RLK known for their role in transpiration efficiency and drought resistance ^35^.

Cross-referencing the 198 candidate genes with RNA-Seq studies on the response of *M. polymorpha ssp. ruderalis* to infection by the pathogen *Phytophthora palmivora* ^36^, and to various abiotic stresses ^11^, allowed to identify 97 of them differentially regulated in at least one tested condition (Table S8). Among them, the coiled-coil NLR (CNL) MpNBS-LRR11 (Mp4g08790), one of the top 3% of genes under balancing selection (Table S3), was found differentially expressed during infection with *P. palmivora* and was detected in GEA for biovariables linked to hot and humid conditions often linked to higher pathogen pressure ^37^. MpNBS-LRR11 may thus play a role in the adaptation of *M. polymorpha* to pathogens.

Most of the *M. polymorpha* ssp. *ruderalis* genes identified through the GEA belong to gene families known to be important for the response to abiotic and biotic stresses in angiosperms (such as peroxidases, LRR-RLK and NLR). This suggests that plant adaptation relies on ancestral mechanisms to adapt to new environment, enhanced by lineage-specific duplications that led to a diversity of paralogs across the different plant lineages.

### Pangenomic variations in *Marchantia*

Intraspecific diversity can be characterized using SNPs but also by focusing on variations in the gene content between accessions. To capture genes showing presence-absence variation pattern, the 128 genomes assembled from Illumina data, the long-read based genomes of the *M. polymorpha* ssp. *ruderalis* CA, BoGa, and Tak-1 accessions, the reference genomes of *M. polymorpha* ssp. *montivagans* and *M. polymorpha* ssp. *polymorpha* ^22^, and *Marchantia paleacea* ^38^ were used to generate the *Marchantia* genus pan-genome (Methods).

To explore the distribution of the gene space annotated for each accession, we clustered the predicted proteomes in hierarchical orthogroups (HOGs) (Table S9). These HOGs represent groups of orthologs, which are either specific to few accessions or shared by most of them (Figure 4a). In particular, from the 25 648 total HOGs, 2 839 were specific to *M. polymorpha* ssp. *ruderalis*, 331 to *M. polymorpha* ssp. *montivagans* and 372 *to M. polymorpha* ssp. *polymorpha* (Table S10). These represent genes that evolved in each subspecies, the higher number for *M. polymorpha* ssp. *ruderalis* being likely due to a larger sampling. We also identified a total of 7292 HOGs shared by accessions from all three subspecies and *Marchantia paleacea* (Table S11). These ‘shared *Marchantia* HOGs’ include genes present across land plants for highly conserved functions, such as auxin biosynthesis and signaling (*i.e.,* ARF, YUC) or the formation of epidermal structures (*i.e., RSL1*). Crossing the list of ‘shared *Marchantia* HOGs’ with published single-cell RNA-Seq analysis ^39^, we found the most significant enrichment for genes belonging to the *Cells surrounding the notch* cluster (Table S11), an expected result given the involvement of these cell populations in structuring the development of *Marchantia*. In addition, genes so far only functionally characterized in *M. polymorpha* Tak-1 clustered in the ‘shared *Marchantia* HOGs’. Among them were genes involved in gemmae (*MpKARAPPO*) and gemmae-cup (*MpGCAM1*) formation, idioblast differentiation (*MpSYP12B*) and air chamber development (*MpNOP1*, Table S11) ^40–42^. The enzyme MpAUS/MpPPO responsible to produce the flavonoid Auronidin possibly involved in stress resistance ^43^ is also found in this list of shared genes. Presence of these genes in the ‘shared *Marchantia* HOGs’ indicates that the molecular mechanisms controlling these traits are likely conserved in the *Marchantia* genus, beyond Tak-1 (Figure 4b).

**Figure 4.**
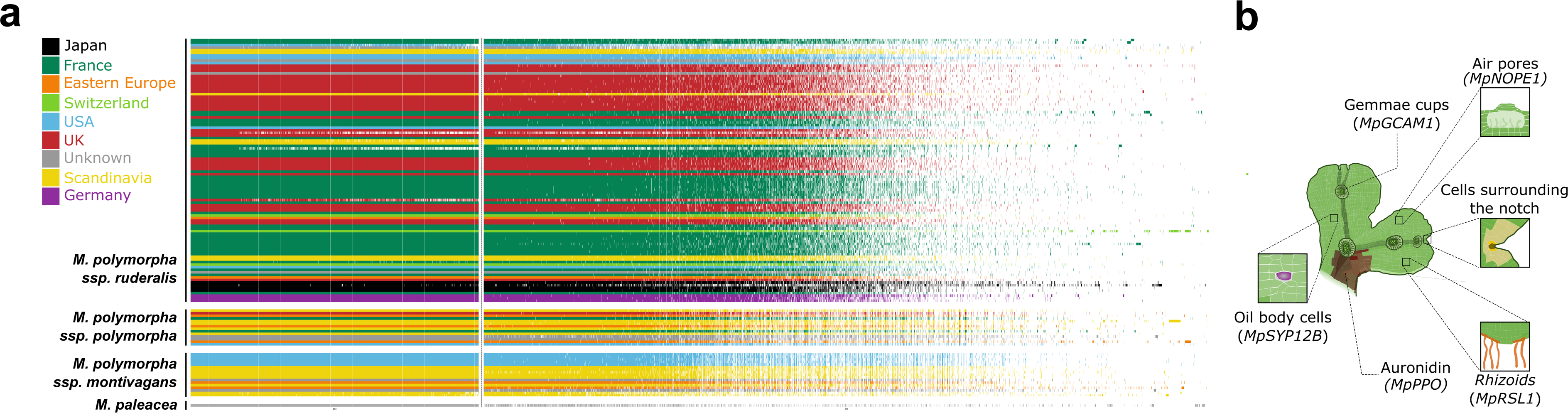
Landscape of gene presence-absence variation in the Marchantia genus pangenome. (a) Hierarchical orthogroups are sorted by their occurrence, with the genes shared by all the *Marchantia* species in the left part (separated from the rest by a dotted line), and the very rare genes in the right part of the matrix. The accessions (in rows) are sorted according to their position in the phylogenetic tree of the accessions generated by the software Orthofinder to infer these HOGs. The colors represent the geographic origin of each accession. (b) Genetic features and traits shared by all species in the *Marchantia* genus.

The *M. polymorpha* ssp. *ruderalis* pangenome was obtained using the gene prediction on 100 ruderalis accessions, the three ruderalis long reads genomes and two outgroups, being the reference genomes from *M. polymorpha* ssp. *polymorpha* and ssp. *montivagans*. A specific Orthofinder analysis was carried out on this sampling to improve the resolution, leading to 28 143 HOGs representing the genic content in the *M. polymorpha* ssp. *ruderalis* (Table S12, Figure 5a). These HOGs were then classified into three categories: core, accessory and cloud genome, depending on the number of accessions with genes in each HOG (Table S13). The core HOGs, present in most *M. polymorpha* ssp. *ruderalis* accessions (Methods), represent 35% of the total pangenome HOG content. Approximately half (49.2%) of the HOGs belong to the accessory compartment of the pangenome (Figure 5b). In this pangenome, the cloud HOGs, present in four or less accessions, are enriched in genes coming from the long-read genomes and should be taken with caution because they may be absent from other accessions for technical reasons (Figure S4).

**Figure 5.**
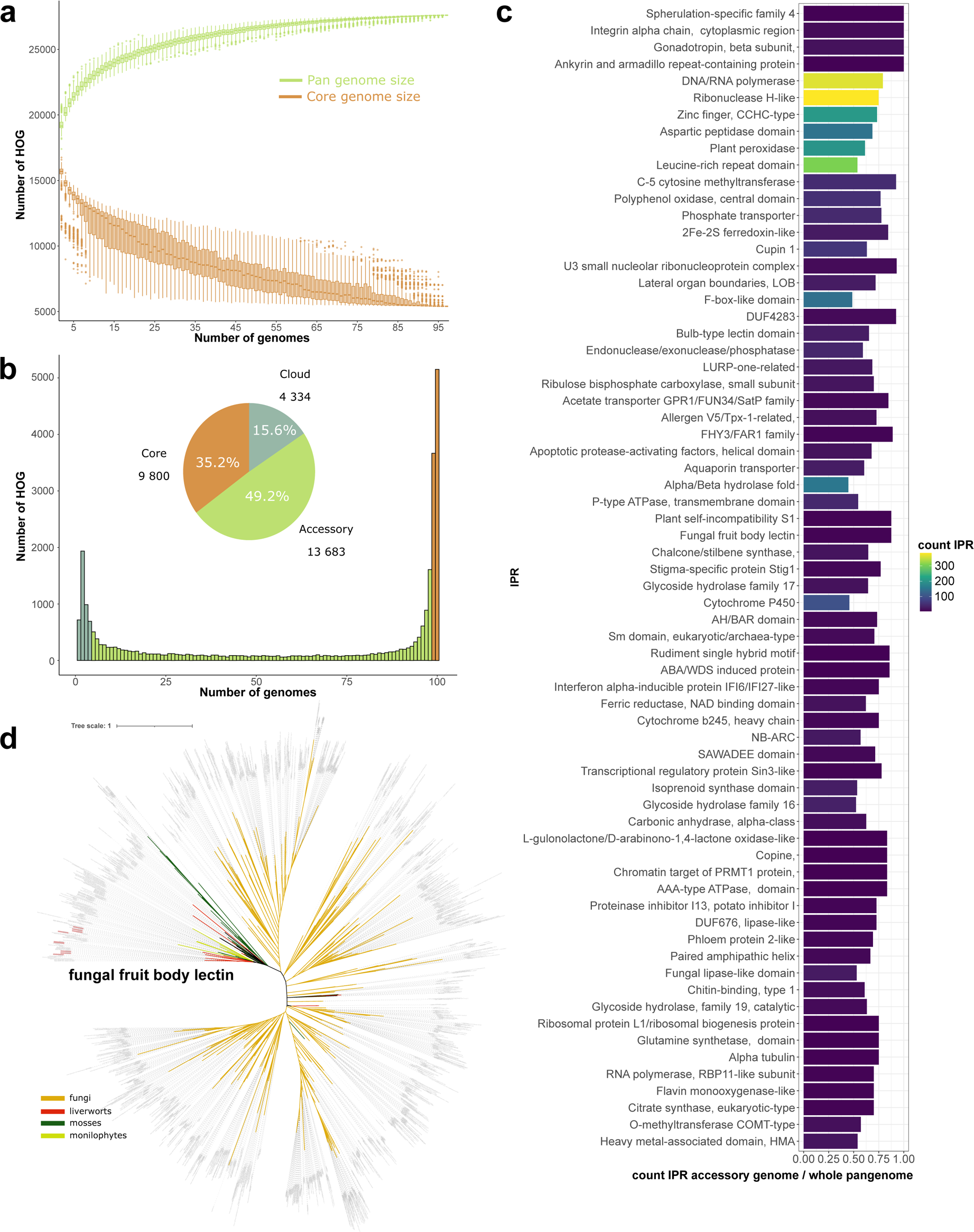
The pangenome of *Marchantia polymorpha ssp ruderalis*. (a) Increase in the pangenome size and decrease in the core genome size when adding accessions to the *Marchantia polymorpha ssp ruderalis* pangenome. This saturation plot indicates a closed pangenome in *Marchantia polymorpha ssp ruderalis*. (b) Composition of the pangenome, regarding the categories of the HOG (core, accessory and rare). The histogram shows the number of HOGs with different frequencies of presence in accessions, and the pie chart shows the proportion of HOG in each category. (c) Barplot representing the IPR significantly enriched in the accessory genome. Redundant IPR and IPR linked with transposable element and viral proteins were discarded to improve readability. (d) Phylogeny of the orthologs of fungal lectin genes present in *Marchantia polymorpha ssp ruderalis*. This tree was computed with the substitution model WAG+R9 and has a log-likelihood of - 121336.9770.

When crossing the lists of core and accessory genes with scRNA-Seq analysis, we found that the expression cluster corresponding to *Cells surrounding the notch* was the most significantly enriched in core *M. polymorpha* ssp. *ruderalis* genes (Table S13), similarly to the *Marchantia* pan-genome. The expression clusters with a significant proportion of accessory genes were the ones linked to *gemma* and *oil body* cells. Among the 283 genes of the *gemma* cell cluster, 83 are accessory and are mainly linked to specialized metabolism, such as chalcone synthases or cytochrome P450. For the *oil body* cell cluster, 52 of the 177 assigned genes are accessory, many of which are also involved in specialized metabolism. This includes some microbial terpene synthase like genes and polyphenol oxidases, probably allowing the on-site synthesis of the oil body metabolites.

The HOG present in the core and accessory compartments of the pangenome were also crossed with RNA-Seq data conducted on Tak-1, as described above for the GEA candidates, and conditions with significantly more core or accessory genes deregulated scrutinized (Table S14). All conditions, excepted the mannitol stress, showed a down-regulation of significantly more core genes than accessory genes (Figure S5). Most conditions also resulted in an overexpression of core genes, but a few conditions induced a higher expression of accessory genes (Figure S5). These conditions include the early stages of the infection by *P. palmivora* (1dpi and 2dpi) and nitrogen deficiency. This indicates that the core genes are more targeted by deregulation than accessory genes during stress responses, but that defined sections of the accessory genome are dedicated to the response to specific environmental constraints.

We performed an InterPro (IPR) domain enrichment on the core and accessory compartments of the *M. polymorpha* ssp. *ruderalis* pangenome. As expected, the core genome is enriched in domains involved in transcription, protein processing, and a variety of processes linked to DNA and RNA (Table S15). These domains are involved in housekeeping functions and most of them are conserved across a wide range of organisms, which is consistent with other pangenomic studies ^44, 45^. The enrichment on the accessory genome included many genes possibly linked to multiple stress responses (peroxidases and isoprenoid synthases), or more specifically to biotic (NLR, chitinases) or abiotic (Aquaporins) stresses (Figure 5c, Table S15).

The *M. polymorpha ssp ruderalis* pangenome expands the role of the accessory genome in the response to biotic and abiotic stresses previously observed in flowering plants ^15^ to a very distant plant lineage, and therefore, potentially, to the entire land plant diversity.

### Horizontal Gene Transfer contributed to plant adaptation to the terrestrial environment

The InterPro domain analysis identified one atypical domain enriched in the accessory genome compartment, the fungal fruit body lectin. Fungal fruit body lectins have been reported in *M. polymorpha* ssp. *ruderalis* Tak-1 and some other liverworts and mosses in 2007 and described as originating from a HGT from a fungal ancestor of the Basidiomycota and Ascomycota clades ^46^. To verify their distribution in the land plants, we searched for orthologs of the nine fungal fruit body lectins present in the reference genome of *M. polymorpha* ssp. *ruderalis* Tak-1, in a database containing genomes from all the main clades of land plants (Table S16) and transcriptomes from the 1KP initiative ^47^. We detected orthologs in liverworts, mosses, but also in *Adiantum nelumboides*, *Alsophila spinulosa* genomes and 21 other fern species transcriptomes (Figure 5d). These lectins had not been identified in ferns so far.

Indeed, these genes are missing from the two previously screened genomes of aquatic ferns ^48^ and may represent a loss following the reversion from a terrestrial to an aquatic environment, as observed for genes related to immunity ^49^ or symbiotic interactions ^50^. This presence of fungal fruit body lectins in terrestrial ferns gives a whole other perspective on the gain of these domains in plants: they might have been transferred from a fungus to the ancestor of land plants and then lost in three clades (hornworts, lycophytes and seed plants) (Figure 5d). Six of the nine fungal fruit body lectin genes present in Tak-1 are differentially expressed in stress conditions, most being up-regulated in response to drought (Table S14).

Collectively, our data indicate that a gene family originating from a fungal HGT before the diversification of land plants contributes to the adaptation to drought in *M. polymorpha* ssp. *ruderalis* while it was independently lost in other lineages, illustrating the different adaptive landscape in land plants after 450 million years of on-land evolution.

## Discussion

Intraspecific diversity in bryophytes had never been explored at a genome-wide scale, resulting in a biased knowledge of plant adaptation mainly restricted to the angiosperms. By gathering a collection of wild-collected accessions of the model liverwort *M. polymorpha* we aimed at providing a resource for the community and at identifying the genetic basis of adaptation in this species. Three main approaches were combined: population genomics-based selection analyses, genome-environment association analyses and a gene-centered pan-genome. By comparing the genes identified through these complementary approaches with knowledge acquired in angiosperms, we were able to identify commonalities in the genetic strategy for adaptation to the environment, but also lineage-specific innovations deployed by

*M. polymorpha*. Two gene families stood out when comparing adaptation in *M. polymorpha* and angiosperms: the class III peroxidases and the NLRs. The class III peroxidases are extremely diversified in angiosperms (75 in *A. thaliana*) as well as in *M. polymorpha* (190 genes), which is surprising considering that liverworts did not experience whole genome duplication like angiosperms or mosses ^51^. In *M. polymorpha* this diversity results from clade-specific tandem duplications, as illustrated by POD128 identified in the GEA analysis (Figure 3). Although the gene family expansion mechanisms are different, the peroxidases are among the genes with the strongest signatures of balancing selection in *M. polymorpha, A. thaliana* and *M. truncatula* correlating with their multiple known roles in development and adaptation in angiosperms ^52^. NLRs were the second class of proteins with a diversity and polymorphism pattern in *M. polymorpha* reminiscent of the one found in angiosperms, clearly associating NLR with adaptation. Among the 35 NLR in the *M. polymorpha* Tak-1 genome, six were found among the top 10% genes under balancing selection and 25 belong to the accessory genome. In angiosperms, NLR diversity has been commonly associated with variability in the ability to deploy immune responses against diverse pathogens. In *M. polymorpha*, although one NLR was found by GEA analysis associated with environmental variables, a direct link with the adaptation to parasites remains to be determined. In the future, conducting Genome-Wide Association Studies in response to pathogens would reveal the genetic basis of resistance in *M. polymorpha* and could confirm the link between NLR and immunity.

While gene families such as peroxidases and NLR associated with adaptation in both bryophytes and angiosperms likely reflect ancestral traits retained since the most recent common ancestor of the land plants, lineage-specific novelties are also expected. These lineage-specific genes can either reflect losses or gains and, if absent from angiosperms, represent potential candidates for crop improvement. Lineage-specific adaptation signatures may have co-evolved with morphological adaptation. In that respect, the importance of the *gemma* and *oil body* cell clusters in the adaptation processes in *M. polymorpha* has been underlined by crossing scRNA-Seq data with the GEA candidates and the accessory genome. Oil bodies are the site of many specialized metabolite storage and synthesis and have already been shown to be involved in *M. polymorpha* resistance to biotic stress ^53^. The farnesyl diphosphate synthase MpIDS1 was found as a GEA candidate associated with temperature and precipitation variables. Since this gene is involved in the synthesis of the farnesyl pyrophosphate, a precursor of sesquiterpenes, it can impact a great diversity of processes related to terpenoids. By looking at the oil body clusters, which mainly express genes related to specialized metabolite synthesis, a clear pattern between the core and accessory genome arise. Core genes (such as IDS1) are more likely to be upstream genes, whereas the accessory genes, such as terpene synthases, likely contribute to final steps of the pathways. The transfer of such terminal genes might represent an option for the production of antimicrobial compounds in crops. An insecticidal protein identified in ferns, Tma12, and absent from other land plants, was successfully transferred to cotton conferring resistance to whitefly ^54, 55^. In ferns, this protein originated from a HGT. Such a case of HGT expanding the abilities to interact with the microbiota is reminiscent of the fungal fruit body lectin proteins, identified in the *M. polymorpha* accessory genome and originating from HGT, previously linked with resistance against insects ^46, 56^. We found that these genes are present in multiple fern genomes and transcriptomes tracing its acquisition to the last common ancestor of land plants. It is therefore possible that other HGT present in bryophytes and lost in angiosperms were acquired by their common ancestor, and that the role of HGT in the terrestrialization process is even more important than what is suspected ^56^. Reinvestigating HGT in the light of the *Marchantia* pangenome and the recently sequenced non-seed plant genomes may allow identifying more of these land plant novelties lost in angiosperms and thus ignored so far.

Our study provides the first genetic and genomic resource to explore intraspecific diversity in bryophytes, expanding the tools available for the model liverwort *Marchantia polymorpha*. Exploring this resource, we identified some genetic basis of adaptation to the environment in this species, revealing similarities with angiosperms together with lineage-specific novelties. *Marchantia* also retains adaptation-related genes inherited from the most recent common ancestor of land plants but lost in angiosperms. This resource provides a reservoir of genetic novelties with the potential to be used for crop improvement.

## Acknowledgments

The authors thank the genotoul bioinformatics platform Toulouse Occitanie (Bioinfo Genotoul, https://doi.org/10.15454/1.5572369328961167E12) for providing computing resources and public contributors of material from the Great British liverwort hunt: Ian Burrow, David G. Hill, The Roberts Family, Peter & Theresa Rooney, Jennie Showers, Anthea Wilson, and an additional 59 anonymous contributors from the British public. Research at the LRSV and LIPMe is supported by the Laboratoires d’Excellence (LABEX)’ TULIP (ANR-10-LABX-41). J.K, C.G, C.L and P-M.D were supported by the project Engineering Nitrogen Symbiosis for Africa (ENSA) currently funded through a grant to the University of Cambridge by the Bill and Melinda Gates Foundation (OPP1172165) and the UK Foreign, Commonwealth and Development Office as Engineering Nitrogen Symbiosis for Africa (OPP1172165). The work (proposal: Award DOI 10.46936/10.25585/60001405) conducted by the U.S. Department of Energy Joint Genome Institute (https://ror.org/04xm1d337), a DOE Office of Science User Facility, is supported by the Office of Science of the U.S. Department of Energy operated under Contract No. DE-AC02-05CH11231. This project received funding from the European Research Council (ERC) under the European Union’s Horizon 2020 research and innovation programme (grant agreement no. 101001675 - ORIGINS) to P.-M.D, from the CNRS to P-M.D and H.P (80|PRIME MicMac), from the Deutsche Forschungsgemeinschaft (ZA, 259/9) to S.Z., from the National Science Foundation to J.M.N. (NSF 1501826), from the Japan Society for the Promotion of Science KAKENHI (JSPS 20K15783), from the Agence Nationale de la Recherche (ANR LEVEL-UP ANR-21-CE20-0010-01), from ZhuJiang (2019ZT08N628) and the National Natural Science Foundation of China (32022006) to S.C, from the URPP Evolution in Action of the University of Zurich, grants of the Swiss National Science Foundation (160004, 131726), the EU’s Horizon 2020 Research and Innovation Program (PlantHUB-No. 722338), the Georges and Antoine Claraz Foundation, and the Forschgungskredit of the University of Zurich (FK-20-089) to P.S.

## Author contributions

P.M-D., J.N., F.R., D.J.H, G.R.L.G, S.S, SLCU Outreach Consortium, P.S., and S.Z. collected and contributed accessions. C.B., M.B., J.K., D.L.M-Z., P-M.D., G.G., N.R., A.T., C.G., P.S., I.D., A.T., S.C., N.R., K.EM., E.A., C.G., S.A., J.W., J.G., W.H., A.M., B.L., A.B. and H.CS carried out experiments. C.B., C.L., M.B., P-M.D, S.C, J.K, H.P. and S.A. analysed data. C.L, M.B., P-M.D, C.J., C.D., H.P., J.L-M., J.S., A.B. and S.C. coordinated experiments. Y.T. developed the website. C.B., M.B., P-M.D wrote the manuscript. F.R., P.S., S.Z. and S.S. Edited the manuscript. M.B. and P-M.D. coordinated the project.

### The SLCU Outreach Consortium

David Hoey (lead), Edwige Moyroud (lead), Alan Wanke, Alessandra Bonfanti, Stefano Gatti, Alexander Summers, Elisabeth Burmeister, Kathy Grube, Andreea Alexa, Nataliia Kuksa, Lauren Gardiner, Martin Balcerowicz, Jemma Salmon, Bryony Yates, Lucie Riglet, Elena Salvi.

## Online method

### Plant material

Plants were collected directly in sampling pots and brought to the lab. After growing individual thallus from each accession in potting soil, gemmae were collected and sterilized as in Delaux et al. 2011 ^57^. Cultures were initiated from single gemma and a single plant conserved for each accession and grown on ½ B5 medium as in Rich et al.^4^ supplemented with 1% sucrose, with 16:8 h day:night cycles at 22 °C under fluorescent illumination with a light intensity of 100 μmol μm−2 s. Note: due to the COVID19 pandemics, a total of 38 accessions, at the time being propagated for the first round of gemmae, were lost.

### BoGa-L5 accession long-read sequencing

#### DNA extraction, library preparation and long read sequencing of BoGa L5

*Marchantia polymorpha* ssp. *ruderalis* ecotype BoGa crossline 5 (BoGa L5) plants were raised on Gamborg B5 medium (Duchefa Biochemie B.V., Netherlands) at room temperature with a day-night cycle of 16 h of light to 8 h of darkness. Male plants were collected at 44 days after germination (DAG) and high molecular weight (HMW) DNA was extracted from 1g of material according to an adapted CTAB-DNA extraction protocol where Chloroform:Isoamyl alcohol was replaced with dichloromethane (https://www.protocols.io/view/plant-dna-extraction-and-preparation-for-ont-seque-kxygxenmkv8j/v1). DNA quality was checked with the Invitrogen Qubit 4 fluorometer (Thermo Fisher Scientific Inc., USA). The DNA was size selected with the short-read eliminator kit (PacBio, USA) and the sequencing libraries were prepared with the ligation sequencing gDNA kit (SQK-LSK109-XL; Oxford Nanopore Technologies (ONT), Oxford, UK). The libraries were sequenced on a GridION Platform using one R9.4.1 and one R10.0 flow cell and bases were called with MinKNOW v1.4.3 software (ONT, Oxford, UK).

#### Short-read sequencing of BoGa L5

A paired-end sequencing library was prepared using the extracted genomic DNA of BoGa and the TruSeq DNA Nano Kit according to the TruSeq DNA Sample Preparation v2 Guide (Illumina, San Diego, USA). The library was sequenced 2×150 bp paired-end on a NovaSeq500 sequencer (Illumina, San Diego, USA). The raw reads were quality trimmed using Trimmomatic v0.39 (Bolger et al., 2014) with parameters ‘LEADING:34 TRAILING:34 SLIDINGWINDOW:4:15 ILLUMINACLIP:2:34:15 MINLEN:100’.

#### Computation of a chromosome-scale BoGa genome assembly

The BoGa long reads were assembled with Canu v2.2^58^ setting ‘genomeSize=280m’. The genome assembly was polished with Racon v1.4.20 ^59^ and minimap v2.22-1101^60^ using the parameters ‘-m 8 -x -6 -g -8 -t 40’, with Medaka v1.4.3 (https://github.com/nanoporetech/medaka) and with Pilon v1.24^61^. Medaka polishing was run one round with the raw ONT reads from R9.4.1 flow cell and parameter ‘r941_min_high_g360’ and one round with the reads from R10.0 flow cell and parameter ‘r103_min_high_g360’. Pilon polishing was done three times using BWA mem v0.7.17^62^ and the trimmed genomic short read data of BoGa L5. The polished contig sequences were anchored to pseudochromosomes based on the Marchantia reference genome sequence assembly Tak v6.1^20^ with RagTag v2.0.1^63^ allowing 100 bp gaps between sequences. For this, chromosome U that represents the female sex chromosome and unplaced scaffolds of the Tak assembly were disregarded and an artificial chromosome 0 was generated from unplaced sequences of the BoGa assembly. The assembly was repeat-masked with RepeatModeler v2.0.2^64^ with parameter ‘-LTRStruct’ and RepeatMasker v4.1.2^65^ with enabled soft-masking option.

The completeness of the BoGa genome assembly was determined with the Benchmarking Universal Single-Copy Orthologs (BUSCO) tool v5.4.4^66^ using the database ‘viridiplantae_odb10’ to 93.1%.

Chloroplast and mitochondrion contig sequences in the BoGa assembly were identified through a blastn search (package v2.11.0+) against the *M. polymorpha* chloroplast (NCBI accession number NC_037507.1) and mitochondrion genome sequence assembly (NCBI accession number NC_037508.1) resulting in one full sequence each. Overlaps were trimmed for the chloroplast contig with Berokka v0.2 (https://github.com/tseemann/berokka) and for the mitochondrion contig through identification of overlapping ends with a blastn search against itself. The contigs were oriented according to the used references with blastn. The gene annotation of the BoGa chloroplast sequence was computed with GeSeq (v2.03)^67^ applying tRNAscan-SE (v2.0.7), Chloe v0.1.0, HMMER v3.3.1 and BLAT ^68^. The gene annotation of the BoGa mitochondrion genome sequence was calculated with GeSeq using tRNAs-can SE (v2.0.7) and BLAT. The results were filtered discarding annotations with HMMER score<15, BLAT score<20 and tRNAscan score<20. As GeSeq outputs overlapping genes resulted from the different prediction programs, only one gene annotation was kept at a sequence region. If the overlapping genes had the same start and end position, the prediction of BLAT was prioritized, followed by HMMER, followed by Chloe and followed by tRNAscan, If the start or end position or both differ and the overlapping genes are named equally, the longest gene annotation was kept. The assembly and all associated data can be accessed at https://doi.org/10.4119/unibi/2982437

### CA accession long-read sequencing

#### Preparation of high molecular weight DNA for sequencing

DNA was isolated from dark treated three-weeks old thalli using QIAGEN Genomic-tips 500/G kit (Cat No./ID: 10262) following the tissue protocol extraction. Briefly, 13.5g of young leaf material were frozen and ground in liquid nitrogen with mortar and pestle. After 3h of lysis at 50°C and one centrifugation step, the DNA was immobilized on the column. After several washing steps, DNA is eluted from the column, then desalted and concentrated by alcohol precipitation. The DNA is resuspended in EB buffer. DNA quality and quantity were assessed respectively using the Nanodrop-one spectrophotometer (Thermo Scientific) and the Qbit 3 Fluorometer using the Qbit dsDNA BR assay (Invitrogen). The size of the DNA was assessed using the FemtoPulse system (Agilent, Santa Clara, CA, USA).

#### Preparation and sequencing of HiFi PacBio library

An HIFI SMRTbell® library was constructed using the SMRTbell® Template Prep kit 2.0 (Pacific Biosciences, Menlo Park, CA, USA) according to PacBio recommendations (PN 101-853-100, version 05). Briefly, HMW DNA was sheared by using Megaruptor 3 system (Diagenode, Liège Science Park, Belgium) to obtain a 20Kb average size. Following an enzymatic treatment on 5 µg of sheared DNA sample for removing single-strand overhangs and DNA damage repair, ligation with overhang adapters to both ends of the targeted double-stranded DNA (dsDNA) molecule was performed to create a closed, single-stranded circular DNA. A nuclease treatment was performed by using SMRTbell® Enzyme Clean-up kit 2.0 (Pacific Biosciences, Menlo Park, CA, USA). Due to the limited quantity of library, no size selection was performed. The size and concentration of the final library were assessed using the FemtoPulse system (Agilent, Santa Clara, CA, USA) and the Qubit Fluorometer and Qubit dsDNA HS reagents Assay kit (Thermo Fisher Scientific, Waltham, MA, USA), respectively. Sequencing primer v2 and Sequel® II DNA Polymerase 2.0 were annealed and bound, respectively to the SMRTbell library. The library was loaded on 2 SMRTcells 8M at an on-plate concentration of 70pM using adaptive loading. Sequencing was performed on the Sequel® II system using the Sequel® II Sequencing kit 2.0, a run movie time of 30 hours with 120 min pre-extension step and Software version 9.0 PacBio) by Gentyane Genomic Platform (INRAE-Clermont-Ferrand, France).

#### Whole genome assembly

The data were assembled using HiFiasm assembler (v16.1^69^). Hifiasm is able to produce a primary assembly and an alternative assembly (incomplete alternative assembly consisting of haplotigs under heterozygous regions. We then filtered the primary assembly from organelles and low quality contigs. Organelles were identified by mapping contigs on NCBI reference assemblies. The closest references (AP025455.1 and AP025456.1) were found by MitoHiFi software^70^. We finally removed 601 contigs from the primary assembly (mitochondrial and chloroplastic contigs identified, high covered (>100X), low covered (<1X) and high %GC (>50%) contigs). All the metrics correspond to the filtered primary assembly used for the next analysis. We obtained 527 046 corrected reads (N50 = 17.6kbp; genome coverage = 23X) that were assembled and filtered, resulting in 109 contigs (N50 = 5.99Mbp; L50 = 12 contigs; GC content = 43.34%).

To assess the completeness and quality of the genome, we used the BUSCO v5.4.4^71^ tool on the viridiplantae odb10 database (n=425^66^). We obtained a 99.1% complete BUSCO score on the assembly.

### Short-read sequencing

#### DNA extraction, library preparation and DNA sequencing (Illumina)

Genomic DNA was extracted for all accessions, either from *in vitro* propagated material or from plants grown in pots. For plants sequenced at BGI-Shenzhen or at a service company, genomic DNA was extracted using a CTAB modified protocol from Healey et al. 2014^72^. Approximately, 0.1g of cleaned young thalli (2-3 weeks) from Marchantia populations were reduced in a fine powder in nitrogen liquid. Because Marchantia tissues contain high amounts of phenolic compounds and other molecules which can interfere with DNA extraction, 1ml of STE (Sucrose 0.25M, Tris 0.1M and EDTA 0.05M) wash buffer was previously added to the fine powder in Eppendorf tube. And then, the mixture was thoroughly vortexed and centrifuged during 5min at 14000 rpm. After centrifugation, the supernatant was removed and the washing of ground tissues with STE buffer was repeted until the supernatant was clear. Once ground tissues were clearly washed, DNA extraction and Nanodrop quality was performed as described Healey et al. 2014 by adjusting volumes to the 0.1g starting powder. Illumina libraries were prepared according to manufacturer instruction. Illumina sequencing was performed using either Illumina sequencing technology (HISEQ2000 and HISEQ4000) or the Illumina NovaSeq 6000 at BGI-Shenzhen or the service company respectively, in both cases using a 2×150 paired-end (PE) configuration. For accessions sequenced at JGI, DNA was extracted using the CTAB/STE method (Shepherd & McLay, 2011). Libraries were prepared following the manufacturer instruction and sequenced on an Illumina NovaSeq 6000 instrument using a 2×150 paired-end (PE) configuration.

All sequenced librairies are available under the PRJNA931118 on the Sequence Read Archive (https://www.ncbi.nlm.nih.gov/sra/)

#### Assembly and cleaning of 129 *M. polymorpha* Illumina sequences

For 129 accessions that had a good enough quality, the short reads were processed, and adaptors removed with Trim Galore! V 0.6.5^73^ with -q 30 option and then assembled with megahit v1.1.3^74^ with default parameters. These assemblies were then cleaned, by discarding the contigs not complying to the following characteristics: GC content inferior to 55, at least one hit but no bacterial one in the top 10 blast vs nt (from february 2023) hits of the contigs, length of the contig superior to 500bp.

#### Prediction and functional annotation of protein coding genes

Genome assemblies were softmasked using Red v2.0^75^ and the structural annotations were conducted using BRAKER-2.1.6 pipeline ^76–85^. BRAKER2 was run with --epmode -- softmasking --gff3 --cores 1 options. In epmode, ProtHint pipeline generates hints for AUGUSTUS training and predicts protein coding genes. The OrthoDB input proteins used by ProtHint is a combination of https://v100.orthodb.org/download/odb10_plants_fasta.tar.gz and proteins from seven species (*Anthoceros agrestis* cv. BONN, *Anthoceros agrestis* cv. OXF, *Anthoceros punctatus* ^86^, *Ceratodon purpureus* strain R40 (NCBI GCA_014871385.1), *Marchantia paleacea* ^38^, *Marchantia polymorpha ssp. ruderalis* TAK1 (https://marchantia.info/download/MpTak_v6.1/), *Physcomitrium patens* ^87^ and *Sphagnum fallax* ^88^). Only the predicted genes partly supported by this protein database were kept using the https://github.com/Gaius-Augustus/BRAKER/blob/report/scripts/predictionAnalysis/selectSupportedSubsets.py script. The long reads genomes of the *M. polymorpha ssp. ruderalis*, *M. polymorpha ssp. polymorpha*, *M. polymorpha ssp. montivagans* ^22^ and *M. paleacea* were reannotated in the same fashion, as well as the two new long read genomes from the accessions CA and BoGa-L5.

The completeness of the predictions in each accession was assessed with BUSCO v5.4.4 against the viridiplantae odb10 (n=425).

The predictions were functionally annotated with interproscan-5.51-85.0 ^89, 90^ with options – iprlookup and –goterms.

#### Mapping and SNP calling of the 135 accessions

Trim Galore! v 0.6.5 ^73^ was used to remove 3’ bases with a quality score lower than 30, trim the Illumina adapters, and only keep the reads longer than 20bp. Bowtie2 version 2.3.5.1 ^91^ was then used to map the reads from the 135 accessions to the reference genome of *M. polymorpha ssp. ruderalis* (Tak-1 accession, MpTak v6.1 genome version, ^92^), without permissiveness for discordant and mixed alignment. The polymorphic variants with a minimum of 4 supporting reads, a minimum base quality of 30, a minimum variant allele frequency of 0.97, and a p-value threshold of 0.01 were then called with VarScan.v2.4.2 ^93, 94^. The resulting VCF was then filtered to discard indels, 2 accessions of low quality, and keep only biallelic sites with information in at least 50% of the accessions, leading to a total of 12 519 663 SNP in the 133 remaining accessions, and 5 414 844 in the 104 *M. polymorpha ssp. ruderalis* accessions.

#### Phylogeny and population structure analysis

A total of 107 934 multiallelic and biallelic SNPs, pruned with the R package SNPRelate (Zheng et al., 2012) for a linkage disequilibrium (LD) < 0.3 on 1 500bp windows, and with missing data < 4% were kept for phylogenetic analysis on the 133 *M. polymorpha* accessions. SNP based phylogenetic tree was inferred with IQ-TREE version 2.1.2^95^ with ModelFinder ^96^) with SH-like approximate likelihood ratio test ^97^ and ultrafast bootstrap ^98^ (with 10 000 replicates). The tree was then visualised using the phytools R package ^99^.

Population structure analysis was performed on 95 non redundant accessions from the *M. polymorpha ssp. ruderalis*. SNPs with a missing frequency < 20%, a minor allele count of 6 accessions and a plink pruning on 10kb windows, a step size of 1 and a LD threshold of 0.5 (458 346 SNPs) were used in fastStructure 1.0^100^. Population numbers (K) from 2 to 7 were tested with 5 cross validation test sets, pointing to an optimal K between 2 and 4.

#### Determination of selection signature on genes

Selection signatures were detected with indicators based on the AFS: Tajima’s *D* ^101^, Fay and Wu’s *H* ^102^ and Zeng’s *E* ^103^. By exploiting the biallelic SNP dataset in the 3 subspecies, the ancestral or derived state of each allele in *M. polymorpha ssp. ruderalis* was determined. For the polymorphic positions in *M. polymorpha ssp. ruderalis* that were fixed the same way in the two other subspecies (allowing missing information in one accession of each subspecies), the *M. polymorpha ssp. ruderalis* allele corresponding to the fixed allele in *M. polymorpha ssp. montivagans* and *M. polymorpha ssp. polymorpha* was considered to be the ancestral allele. This led to a reduced dataset of 1 344 013 unfolded SNPs that was used to calculate the *H* and *E* indicators. We used the folded SNPs dataset (5 414 844 SNPs) to calculate Tajima’s *D*. The calculation of the *D*, *H* and *E* statistics was performed with a custom R script (https://figshare.com/s/715a36bc7585d46d0279): *θ*_*W*_, *θ*_*π*_ and *θ*_*L*_ were calculated for each SNP, with 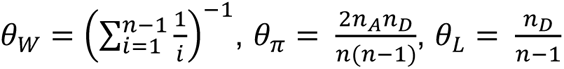 (with *n*_*A*_ and *n*_*D*_ being the copy number of the ancestral and derived allele, respectively, in the case of *H* and *E* calculation), which allowed the correction of the values for the missing data present at each site. The *D* = 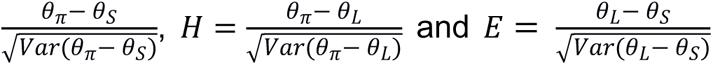 values obtained for each site were then averaged over the SNPs present in each of the 18 620 gene models, allowing to determine the global selective pressure acting on each gene. Genes with pronounced selection signatures were determined based on a dataset of 18 140 genes with at least 4 folded SNPs to calculate the Tajima’s *D* statistics. Based on this, we selected the top 10% of genes (i.e., 1 814) with the highest values of Tajima’s *D* as a gene set with the most pronounced signature of balancing selection. Among the remaining 16 326 genes, based on ancestral/derived SNP dataset (1 344 013 SNPs) we selected 11 932 genes for which the ancestral/derived alleles identification could be inferred for at least 50% of their SNPs. This gene set allowed to select 777 genes with marked signature of background selection based on a Zeng’s *E* values < 10% genome-wide (18 140 genes) quantile value. Finally, the same gene set allowed to select 1,374 genes with marked signatures of soft or hard selective sweep based on Fay and Wu’s *H* values < 10% or *E* values > 90% genome-wide (18 140 genes) quantile value. Hence, 3 965 genes were considered has having pronounced selection signatures based on three neutrality test statistics *D*, *H* and *E*.

The calculation of *D*, *H*, *E* statistics and the selection of genes with pronounced selection signatures was performed in the same way for the angiosperm models species *A. thaliana* and *M. truncatula.* For *A. thaliana*, the SNP dataset from the 1 001 genomes consortium (https://1001genomes.org/data/GMI-MPI/releases/v3.1/; ^104^) was used, coupled with mapping and calling of the *A. thaliana* SNPs on sequencing data from 30 *A. halleri* (PRJNA592307) and 29 *A. lyrata* (PRJNA357372) accessions on the *Arabidopsis* TAIR10 reference genome. Read from the two outgroups were processed with Trim Galore! similarly to what was done on *M. polymorpha*, and then mapped with Bowtie2 version 2.3.5.1 with permissiveness for discordant and mixed alignment, and with an alignment score lower than the minimal score threshold −1.5 + −1.5*L to allow mapping of reads from distant species. The polymorphic variants present in *A. thaliana* were called with VarScan.v2.4.2 in the two other subspecies, with a minimum of 4 supporting reads, a minimum base quality of 30, a minimum variant allele frequency of 0.97, and a p-value threshold of 0.01, leading to 9 500 949 sites for the *ssp. halleri* and 9 206 703 for the *ssp. lyrata*. The unfolded *A. thaliana* sites were the ones bearing the same allele in 100% and 90 % of the accessions from ssp. *halleri* and *ssp. lyrata*, respectively, and with less than 30% of missing values. This led to 3 657 821 SNPs with ancestral allele information, on which the *H* and *E* were calculated. Tajima’s *D* was calculated on the 11 458 975 SNPs from the 1 135 accessions of the 1 001 genomes consortium. For *M. truncatula*, the SNP dataset based on mapping of 317 accessions (among which 285 are from the *M. truncatula* species) on the version 5 of the A17 genome was used (https://data.legumeinfo.org/Medicago/truncatula/diversity/A17.gnm5.div.Epstein_Burghardt_ <u>2022/;</u> ^105, 106^). The 50 885 956 biallelic SNP with minimum base quality of 30, no half calls end maximum 50% of missing data were used to calculate Tajima’s *D*. The SNP calling of 8 accessions from various other species phylogenetically close to the *truncatula* species (species *turbinata, italica, doliata, littoralis, turbinata, tricycla and soleirolii*), performed by Epstein and collaborators, was taken as the outgroup dataset. The ancestral allele was determined for sites that were fixed the same way in all 8 outgroup accessions, with a tolerance for missing data in one accession only, leading to a dataset of 12 129 573 unfolded *M. truncatula* SNPs, on which *H* and *E* were calculated. The values of the *D*, *H* and *E* statistics calculated for *A. thaliana* and *M. truncatula* SNPs were then averaged over their gene models.

#### Estimation of recombination rates and of LD decay

The genome wide landscape of recombination was determined based on the SNP data, using ReLERNN v1.0.0 ^107^. For the simulation phase, the assumed mutation rate was 10^-8^, and the assumed generation time to be 1 year. The other phases (train, predict and BScorrect) were run with default parameters. The LD decay in *M. polymorpha ssp. ruderalis* was determined on the dataset of 5 414 844 SNPs called on the 104 *ssp. ruderalis* accessions, using PopLDdecay ^108^ with a MAF of 0.05 and a maximum pairwise SNP distance of 40 kb. The LD decay was also computed on each chromosome separately.

#### Genome-Environment Association analyses

GEA analyses were carried out to identify genomic regions whose SNP alleles displayed statistical association with the bioclimatic variables extracted from the Worldclim2 database (https://www.worldclim.org/, ^109^). We performed analyses on each of the 19 bioclimatic variables (BIO1 to BIO19), on monthly average precipitations, solar radiation (kJ m-2 day-1) and water vapor pressure (kPa) and on elevation (meters above sea). To integrate the correlations among bioclimatic variables, we performed a Principal Component Analysis (PCA) separately of temperatures (BIO1 to BIO11) and precipitation (BIO12 to BIO19) variables, monthly precipitations, solar radiation (kJ m-2 day-1) and water vapor pressure (kPa). The first 3 principal components were then used for the GEA analyses. GEA analyses were performed using the mixed linear model implemented in GEMMA software (v 0.98.1 ^110^). We used a dataset of 2 155 434 SNPs with minor allele frequencies of 0.05 and maximum 10% missing data, on a set of 96 accessions of *M. polymorpha ssp. ruderalis* for which climatic data were available. To estimate the SNPs effects and their significance, the model used a centered kinship matrix as a covariable with random effect, and a Wald test. SNP *P*-values from GEMMA were processed using a local score approach ^111, 112^ to help detect robust loci across analyses. The local score is a cumulative score that takes advantage of local linkage disequilibrium among SNPs. This score, defined as the maximum of the Lindley process over a SNP sequence (i.e., a chromosome), was calculated using a tuning parameter value of ξ=2, as suggested by simulation results ^111^. Chromosome-specific significance thresholds (ɑ =5%) were estimated using a resampling approach. The R scripts used to compute the local score and significance thresholds are available at https://forge-dga.jouy.inra.fr/projects/local-score/documents.

#### Orthofinder analysis on representative sample of land plants

To compare genes across evolutionarily distant species *A. thaliana, M. truncatula and M. polymorpha* orthogroups have been reconstructed using OrthoFinder v2.5.2 ^113^. To ensure an accurate reconstruction of orthogroups, 36 species covering the main lineages of land plants were added to the above-mentioned species. A first round of OrthoFinder was performed and a careful inspection of the estimated species has been conducted to ensure the absence of errors compared to the phylogeny of land plants ^47, 114^. A consistent tree has been reconstructed and the tree has been used for a second run of OrthoFinder with the option “msa”, enabling to reconstruct the orthogroups using alignment and phylogenetic reconstruction methods.

#### Gene-centered pangenome construction

The presence-absence variation of genes in *M. polymorpha*, and in the subset of *M. polymorpha ssp. ruderalis* accessions, was determined based on orthogroup inference. The proteins predicted on the short reads and long reads assemblies were pooled together and Orthofinder was run independently on the dataset of 104 genomes for the *ssp. ruderalis* pangenome, and on the dataset of 134 genomes for the Marchantia complex pangenome. The Orthofinder analyses were run a first time with default parameters, the resulting species tree was then re-rooted (on *Marchantia paleacea* for the Marchantia complex pangenome, and on the *ssp. montivagans* reference genome for the *ssp. ruderalis* pangenome) and a second Orthofinder run forced on this tree topology was performed, with the multiple sequence alignment parameter. To remove any remaining contamination, all the proteins used in this orthogroup inference were blasted against the non-redundant protein database from the NCBI (NR version from 2023-02-20) using diamond v2.0.8^115^ with an e-value threshold of 10^-3^, a maximum of 10 target sequences and a block size of 4. Hierarchical orthogroups (HOG) with 75% of viridiplantae diamond hits over all genes were considered as reliable, the other ones were discarded. This cleaning led to 25 648 HOGs in the Marchantia complex pangenome (initially 33 418 HOG), and 28 143 HOG in the *ssp. ruderalis* pangenome (initially 35 004 HOG). Two accessions were removed from the ensuing analyses because they had less than 10 000 genes assigned to HOG. Core orthogroups were determined in each subspecies. For *ssp. montivagans* orthogroups containing 15 out of the total 17 accessions were considered core (because two accessions from this subspecies were absent from many core orthogroups), when for *ssp. polymorpha* core orthogroups had to contain all accession. For the *ssp. ruderalis* the core orthogroups were defined as orthogroups containing all accessions, with tolerance for six specific accessions with lower quality, meaning that some or all of them could be absent from the HOG and the HOG would still be considered as core in the subspecies. The list of shared orthogroups among the Marchantia complex was determined by crossing those core orthogroup lists in each subspecies with the list of orthogroups present in *M. paleacea*. HOGs were considered specific to a subspecies if they contained at least one accession of the subspecies and were absent from all accessions of other subspecies. For the *M. polymorpha ssp. ruderalis* pangenome, the core genome was determined the same way as in the Marchantia complex pangenome (shared by all accessions with the tolerance margin for the 6 lower quality accessions) but on the orthogroups generated by the orthofinder run on the 104 genomes. The accessory genome was constituted of the remaining HOG that are present in more than four accessions, and the cloud genome comprises HOG present in four accession or less.

#### Pangenome saturation

The *ssp. ruderalis* pangenome and core genome saturation was studied by determining the number of HOG present (whole pangenome) and the number of HOG shared between the accessions (core genome) for a given number of accessions. For each given number of accessions, 500 random samplings were conducted. The corresponding numbers of HOG present in the sample and shared between accessions were plotted separately as boxplots.

#### Pangenome IPR enrichment

Enrichment analyses were performed on the compartments determined on the *M. polymorpha ssp. ruderalis* pangenome, comparing the IPR domains present in the core and accessory genome to the content of the whole pangenome. The presence of multiple genes for an accession in the same HOG was accounted for by multiplying the IPR count of each HOG by the median number of gene per accession in the HOG. The IPR content of each compartment was then compared to the IPR content of the whole pangenome with a hypergeometric test, IPR present less than 5 times in the whole pangenome were filtered out, and multiple testing correction was computed on the remaining IPRs (Benjamini Hochberg FDR). The IPR with a corrected p-value less than 0.05 were considered as significant.

#### Phylogenetic analysis

The orthologs of some GEA candidates (like ABC1K7) were determined by BLASTp+ v2.12.0 ^116^ (maximum of 4000 target sequences and e-value of 10^-30^) against a database of 37 Streptophytes (Table S10). The resulting sequences were aligned with muscle5 v5.1 ^117^ and trimmed with trimAl v1.4 ^118^ to discard positions with more than 60% of gaps. The phylogenetic tree was computed with IQ-TREE v2.1.2 and the best-fitting evolutionary model selected using modelFinder according to the Bayesian Information Criteria. Branch support was estimated with 10 000 replicates of both SH-like approximate likelihood ratio and ultrafast bootstrap. The phylogenetic analysis of *Marchantia polymorpha* fungal lectins was performed by blasting (BLASTp+ v2.12.0) their protein sequence against transcriptomes of the 1KP initiative ^47^, a database of fungal genomes from MycoCosm (^119^ last time consulted 02/2019), a database of algal genomes and a database of plant genomes (table S19), each with a e-value of 10^-5^ and maximum 2 000 target sequences. Muscle5 was used to align the sequences, trimAl to discard positions with more than 60% of gaps and IQ-TREE2 to compute the tree, in the same way as the previous phylogenies.

#### Data availability

The SNP data mapped on the Tak-1 assembly (VCF format) and genome assemblies (FASTA) as well as their annotations (protein FASTA and GFF) for the 133 accessions sequenced in this paper are available at MarpolBase (https://marchantia.info/). The *Marchantia polymorpha ruderalis* accession CA annotated genome assembly and raw sequenced data are also available in NCBI under BioProject PRJNA1021402. The BoGa genome assembly is available under https://doi.org/10.4119/unibi/2982437

