## Supplementary material for "The *Marchantia* pangenome reveals ancient mechanisms of plant adaptation to the environment": Figures S1 - S5

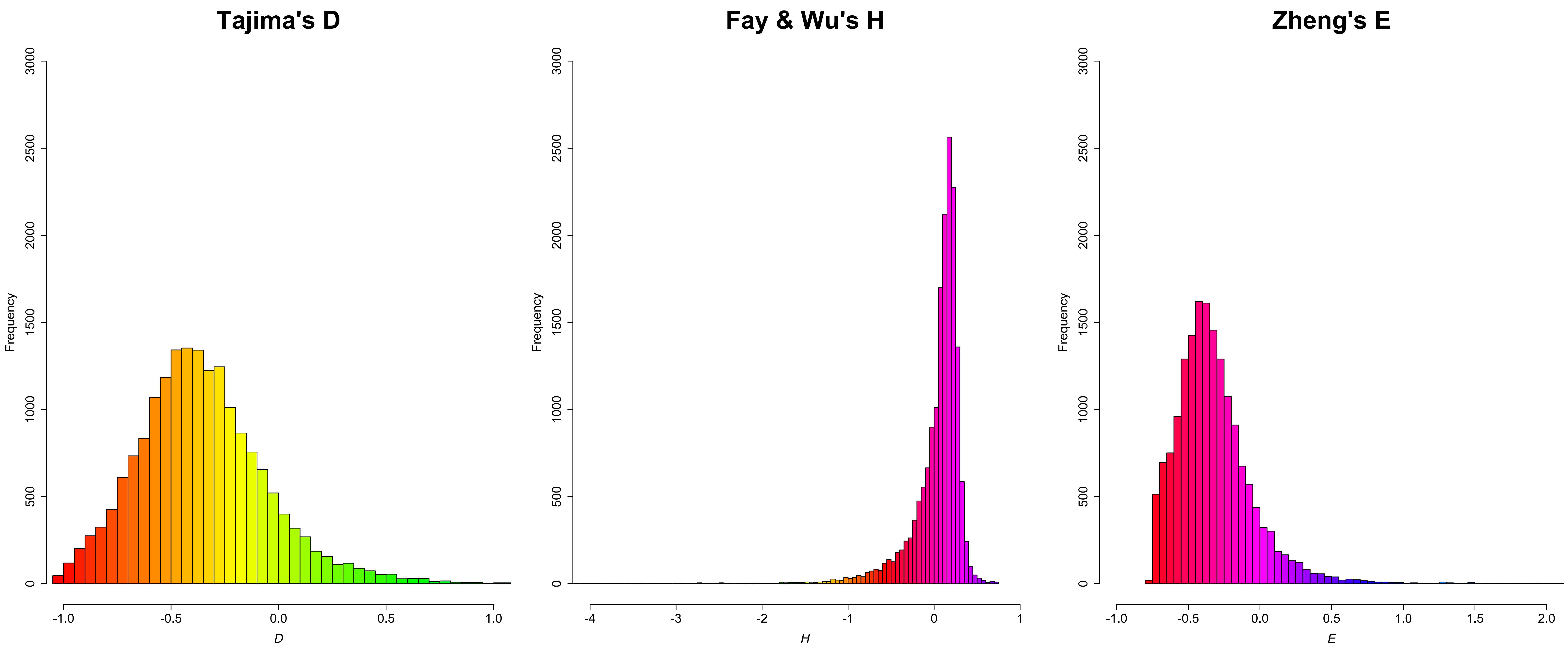


**Supplementary Figure 1 Genome wide gene-based distributions of the D (18 140 genes), H and E (17 027 genes) statistics** based on a dataset of 5 414 844 SNPs for D and 1 344 013 SNPs for H and E, in a collection of 104 accessions of *Marchantia polymorpha* ssp. *ruderalis*.

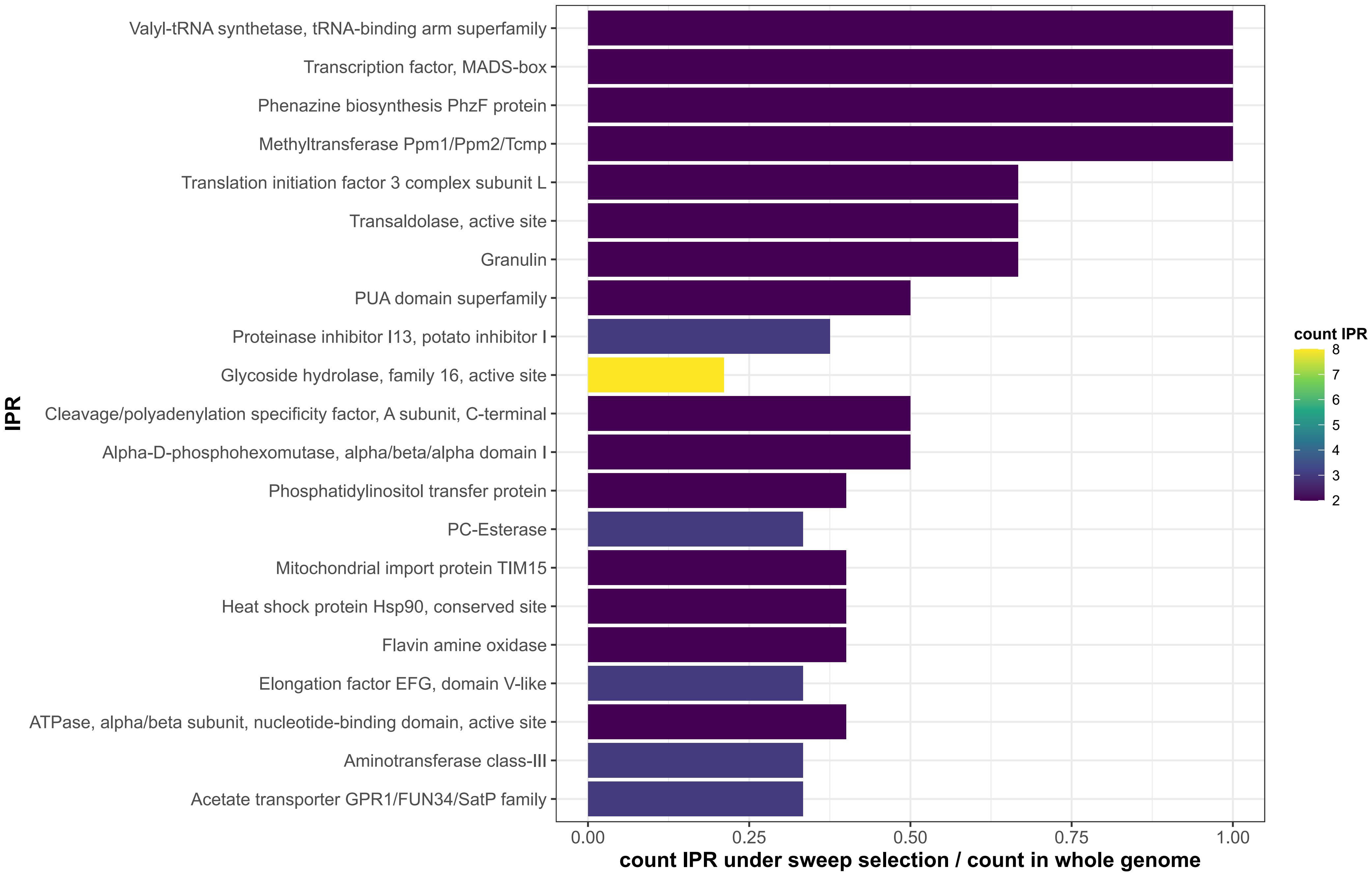


**Supplementary Figure 2 Functional enrichment analysis of a list of 1374 genes putatively under hard or soft selective sweep** (top 10% genes of Fay & Wu's H negative values or Zheng's E positive values) in Marchantia polymorpha ssp ruderalis. IPR terms are ordered from top to bottom according to decreasing significance (FDR q-value < 0.05), together with some associated known gene names.

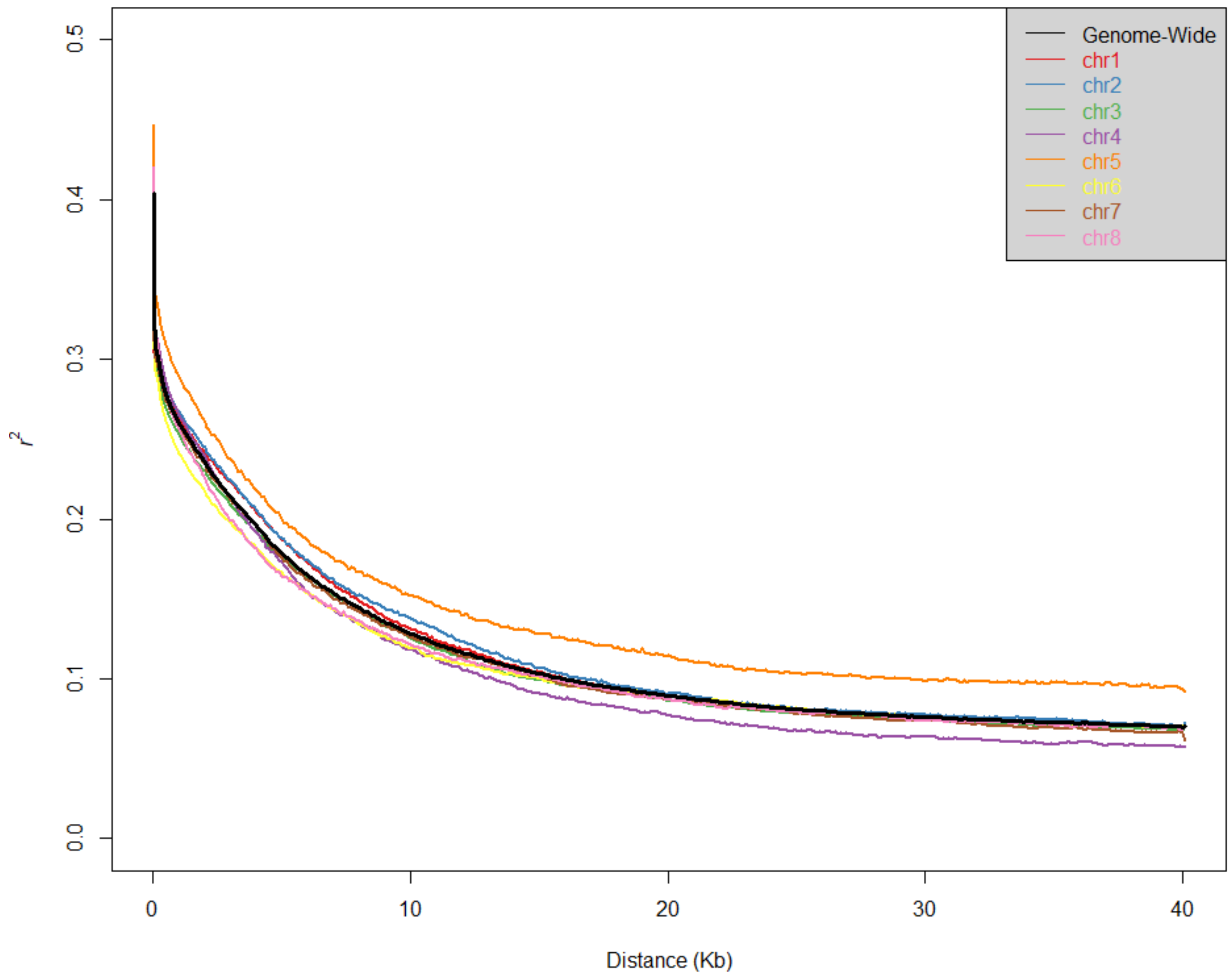


**Supplementary Figure 3 Genome-wide and chromosomal decay of linkage disequilibrium (LD) in a collection of 104 accessions of Marchantia polymorpha ssp ruderalis.** Estimation of LD decay (mean r2 between two SNPs within a given distance bin) were performed using the software POPLDDECAY (ZHANG et al. 2019), based on subsampling of 4 797 046 genome-wide SNP (corresponding to 842 281, 730 749, 785 723, 687 426, 488 279, 680 879, 581 709 and 493 788 SNPs on chromosome 1 to 8), with a MAF threshold of 5%.

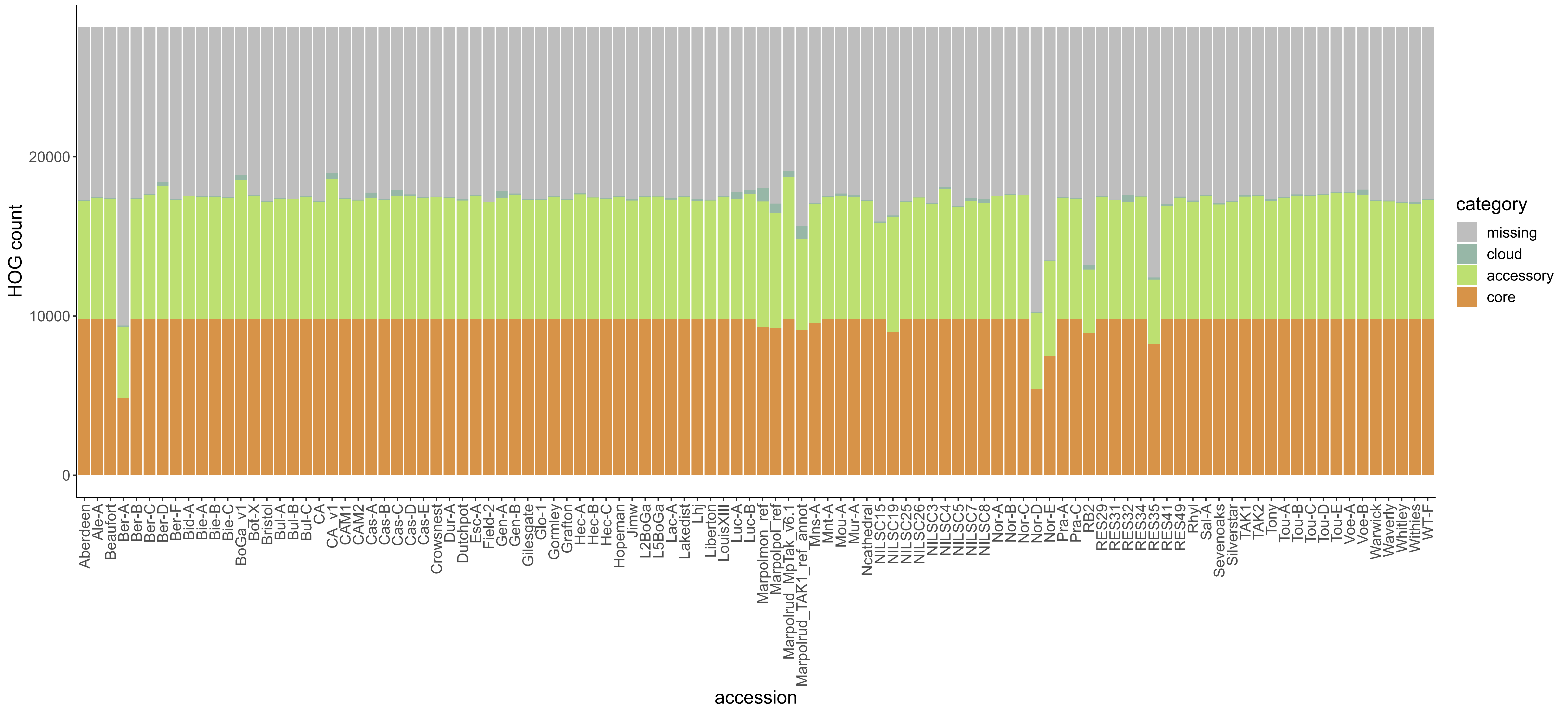


**Supplementary Figure 4 Distribution of the core, accessory and cloud compartment in each of the 102 Marchantia polymorpha ssp ruderalis sequences** (corresponding to 98 accessions with a long and a short read sequencing for CA and BoGa, and a short read sequencing and two annotations of the long reads genome for the reference accession TAK1). The two outgroups, being the reference accessions for the ssp. montivagans and ssp. polymorpha, are also represented.

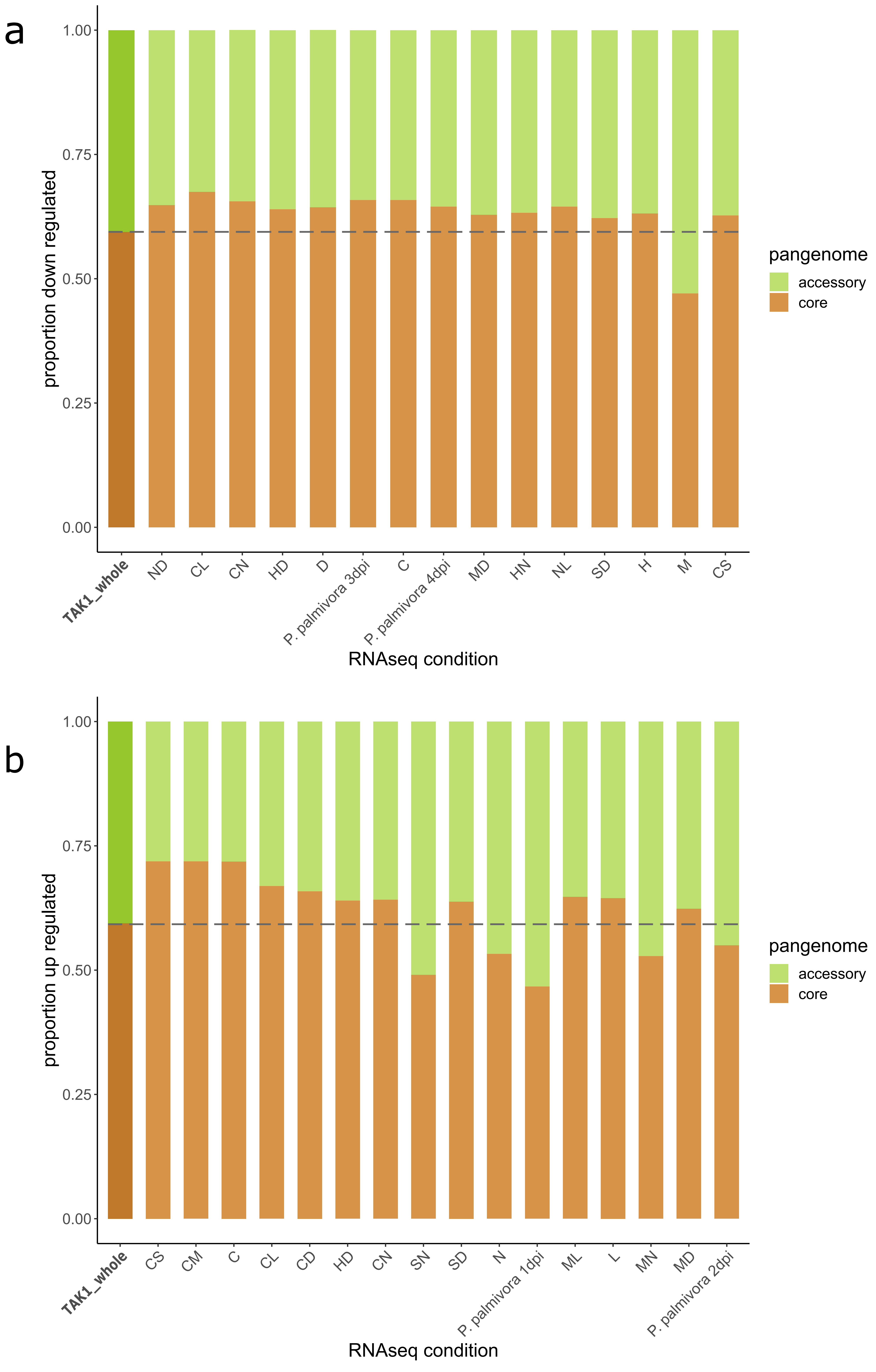


**Supplementary Figure 5 Proportion of genes from the reference genome (Tak-1 v6.1) down (a) or up (b) regulated in stress conditions.** The represented conditions are the ones for which the proportion of core and accessory genes deregulated is significantly different from the proportions of core and accessory genes in the reference genome (represented by the first bar and the dotted line). Signification of the letters for the stress conditions are the following: N nitrogen deficiency, D dark, C cold, H heat, M mannitol, L light, S salt.
